# A Method to Correct Internal Standard Normalization Errors in Sequencing Data

**DOI:** 10.64898/2026.11.23.768725

**Authors:** Robin Graham, Mikhail Tikhonov

## Abstract

Measuring the absolute abundances of microbial communities using only sequencing requires the introduction of an internal DNA standard to all samples. However, any error in the measurement of the internal standard becomes a systematic error propagated to all taxa in a sample. In this work we present a method utilizing random combinations of low-count sequences to detect and correct these systematic errors. When applied to sequencing data from *Lee et al. (2025)*, the method reduced replicate-replicate variance in 66% of samples and did not increase replicate-replicate variance in the remaining samples. This is consistent with a systematic error that is not present in all samples but can be significant when it occurs. Additionally, we show that prior expectations based on the experiment can be incorporated into the method to improve corrections further.

## Introduction

One of the most common classes of measurements in ecology is compositional; what taxa are present, in what amount, and how this composition varies. The standard method for measuring the composition of natural microbial communities has been 16S rRNA genetic sequencing [1]. To compare between samples, sequencing data needs to be normalized to remove sample specific technical biases. Normalizing by the total number of counts converts counts to relative abundances. While historically used, relative abundances introduce spurious correlations between taxa, are unable to detect shifts in total biomass, and can lead to significantly different results when compared to measures of absolute abundance [2].

One technique to measure abundances in absolute units is to introduce a consistent amount of DNA foreign to the sample immediately before DNA extraction [2, 3]. The sequenced counts attributed to this internal standard (ISD) in ideal circumstances can be understood as representing the same quantity of DNA in all samples independent of other technical variations between samples. ISD’s of this type are commonly referred to as ‘spike-ins.’ Taxon counts normalized by a spike-in can similarly be thought of as a proxy for taxon absolute abundance, but counts normalized by a spike-in are still only an indirect measure of absolute abundance because individual sequences exhibit unique amplification biases amongst other biological or chemical differences.

However, any error in spike-in measurement creates a systematic error that is passed on to every absolute abundance, resulting in a over- or under-enrichment of all taxa in a sample(**Fig. 1**). In this work we introduce a method for identifying and correcting spike-in errors. We show that randomly assembled groups of low-count sequences can be used to detect spike-in errors. Utilizing low-count sequences keeps idiosyncratic noise (or actual biological variations) small, while combining them into a large group increases the signal of any systematic error. We verify our method on wild soil microbiome sequencing data collected by *Lee et al. (2025)* [4]. Finally, we demonstrate that prior beliefs about the data can be used in our method to improve spike-in error corrections.

**Figure 1.**
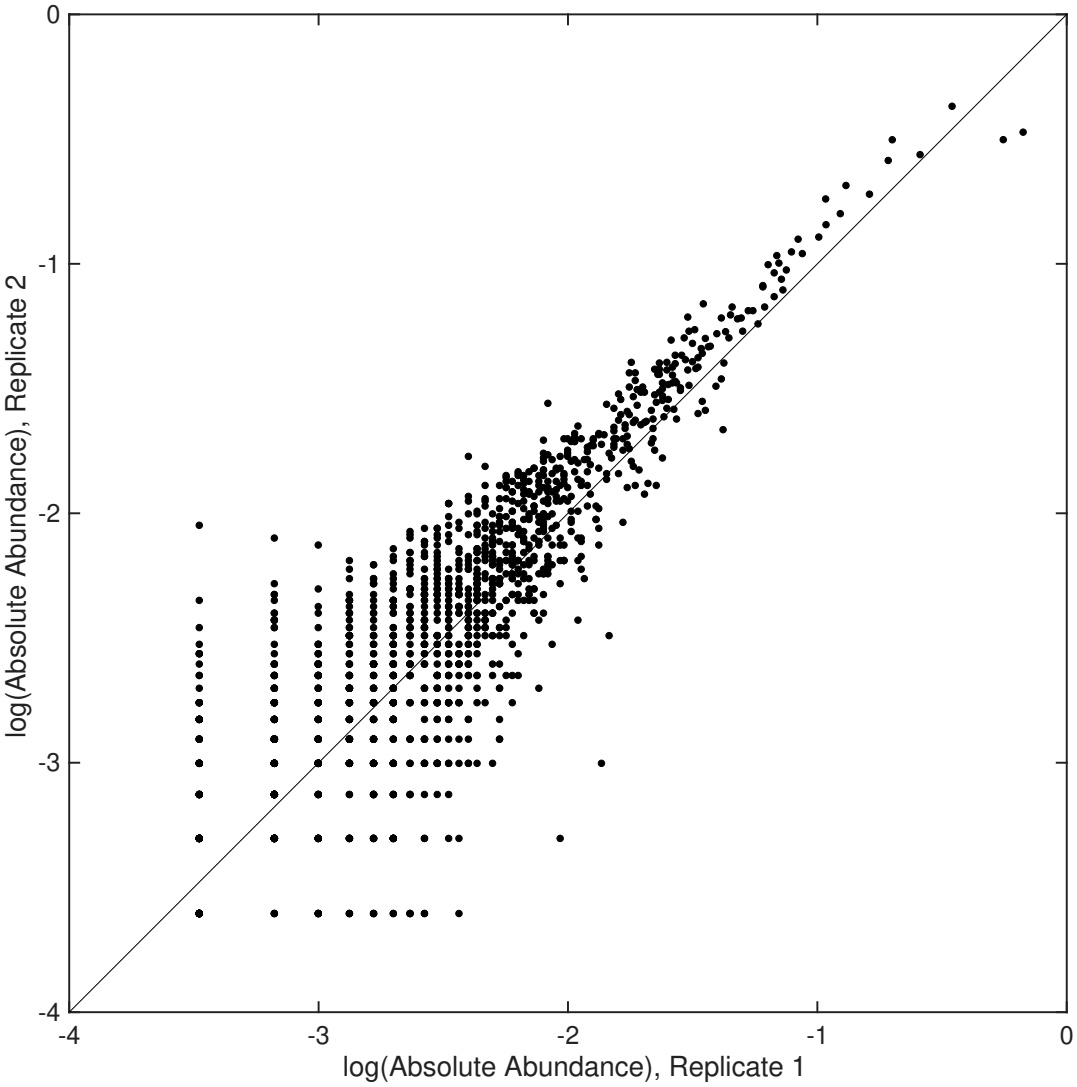
Internal standards can introduce systematic errors. The log absolute abundance of sequences measured in two technical replicates from *Lee et al*. (specifically samples A76 and A77 in Soil 5). In an ideal sample, there will be stochastic differences between technical replicates that can arise from biological variations and measurement noise. An internal standard measurement error will result in a systematic over- or under-abundance of sequences when compared to other technical replicates. The absolute magnitude of this error is proportional to a sequence’s absolute abundance (**Eq. 3**), so the error is most clear in more abundant sequences. However, there are still cases where large sequences do not follow the overall systematic trend due to real biology or idiosyncratic noise. Combining random low-count sequences into a larger collection of counts displays systematic errors similar to high-count sequences, but without sources for similar large variability.

## Results

### Random groupings of low-count sequences can be used to correct spike-in errors

Of primary concern when generating corrections to spike-in errors (instances when spike-in counts are over- or under-estimated) is that the corrections mistakenly correct real biological signal, idiosyncratic noise, or technical biases. Our method utilizes random groupings of sequences to reduce the risk that a biological signal correlated with phylogeny is mistaken for a spike-in error. Using groups of low-count sequences exploits the portfolio effect, where the variance of the group becomes dependent on the average variance of individual group members; the choice to use low-count sequences constrains that average variance to be small. Finally, we only include sequences that are present in all technical replicates to reduce technical biases from varying sequencing depths. For a full analysis of the underlying noise model, see **Methods**.

We verify our method using experimental 16S sequencing data from wild soil samples collected and sequenced in *Lee et al. (2025)*. If a correction is truly correcting a systematic spike-in error then it should reduce the replicate-replicate variance of any random group of low-count sequences in that sample. To test our method, we generate corrections for every replicate of a sample using a subset of the available low-count sequences and apply those corrections to the remaining low-count sequences that were not used in training. Reductions in replicate-replicate variance are indicative of an accurate spike-in correction, while samples that show no change in variance indicate the absence of a detectable spike-in error. If a sample’s replicate-replicate variance increased it would be a sign that the correction was overfitting to sequence or replicate specific variations. We find that our method decreased replicate-replicate variance in the testing set in 66% of samples, and never increased it (**Fig. 2A**). To test the sensitivity of our corrections to idiosyncratic sequence variation, we split the data in half and compared the corrections obtained from each half. We find that independent estimates of the spike-in error correction agree (**Fig. 2B**). While most corrections are small (typically less than 5%), a subset are large, indicative of a systematic error that can significantly skew some samples but is not universal (**Fig. 2A inset**).

**Figure 2.**
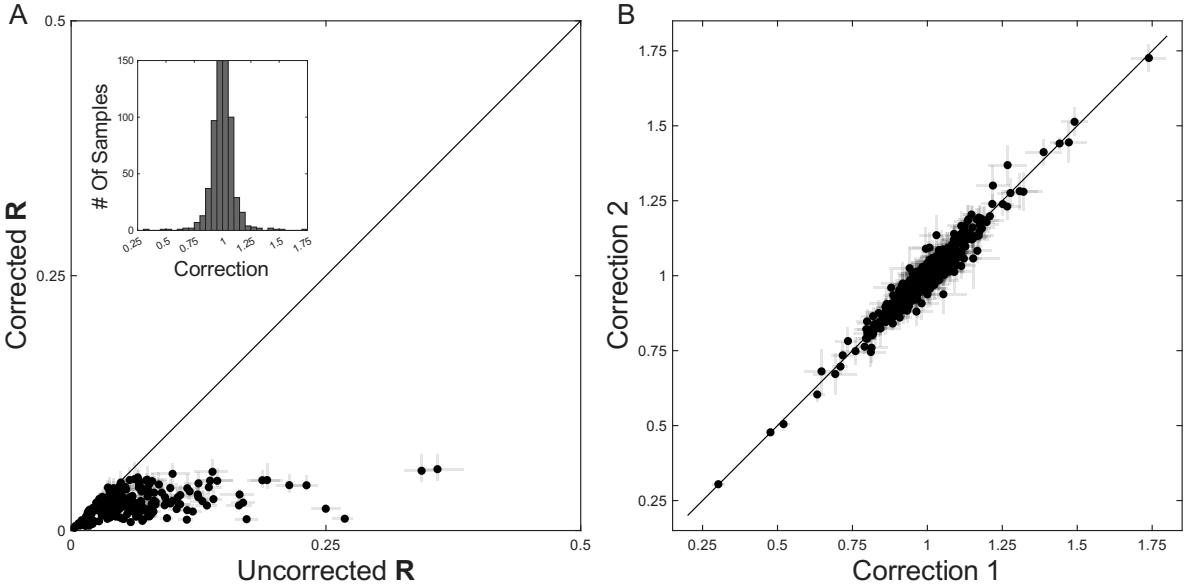
Correcting spike-in errors to reduce variance in groups of small ASVs reduces variance in testing sets. **A:** To test the efficacy of corrections for spike-in errors, corrections generated from a training set of 80% of the data were applied to an independent testing set of the remaining 20%. **R** is the standard deviation across experimental replicates of the absolute abundance of a coarsened group of small ASVs. Each point represents one experimental condition with three technical replicates (except for two conditions with only two technical replicates); points lying below the 1:1 line indicate that corrections reduced variability in the testing set. Error bars show the IQR of **R** across 100 splits of training and testing data. **A, inset:** The distribution of all corrections generated for the data from *Lee et al. (2025)*. **B:** To estimate the quality of corrections, the data for each sample was split in half and two corrections were generated and compared. Each point represents a single experimental replicate with two corrections generated for it different halves of the data; points lying on the 1:1 line were not significantly impacted by being generated from different halves of the training data. Error bars show the average difference between corrections generated from different halves of each already split half of the training data, averaged across 100 splits of each training set split. These repeated splits reduce the usable sequences significantly, so error bars in **B** are larger than for corrections generated from the entire training data. When corrections are generated on the entire dataset, 99% of samples have correction error bars within 5% (**Fig. S3**).

These results can be contrasted with the same method but with large family-level groupings or groupings of high-count sequences instead of low-count sequences. While in some samples these alternative groupings can accurately be used to correct spike-in errors, in others their variance is too dominated by idiosyncratic noise or biological signal and using them leads to significantly increased replicate-replicate variance (**Fig. S1A-S2A**). Corrections using high-count sequences or large family groupings fail to produce consistent corrections when the data is split (**Fig. S1B-S2B**).

### Prior expectations can be incorporated to improve corrections

The baseline method reduces deviations from the average value of a random group of low-count sequences across technical replicates. Using the average abundance is based on a conservative prior that technical replicates will on average be accurate, but a smarter prior could be used if additional information is known. In the case of *Lee et al. (2025)*, pH was the primary perturbation applied to samples. One example of a more informed prior is the expectation that sequence abundance varies smoothly with pH; this can be incorporated into the method by replacing the average across replicates with the average across replicates in the sample together with the replicates in the two samples at neighboring pH values. To evaluate these corrections we compare the abundance of each held-out group of low-count sequences in three sample replicates to its average across nine “sibling” replicates: the three replicates in the same sample and three replicates from each of the two neighboring pH conditions. The pH-smoothness prior brings held-out testing data closer to this sibling average than the standard correction does (**Fig. 3**), indicating that this additional information improves corrections.

**Figure 3.**
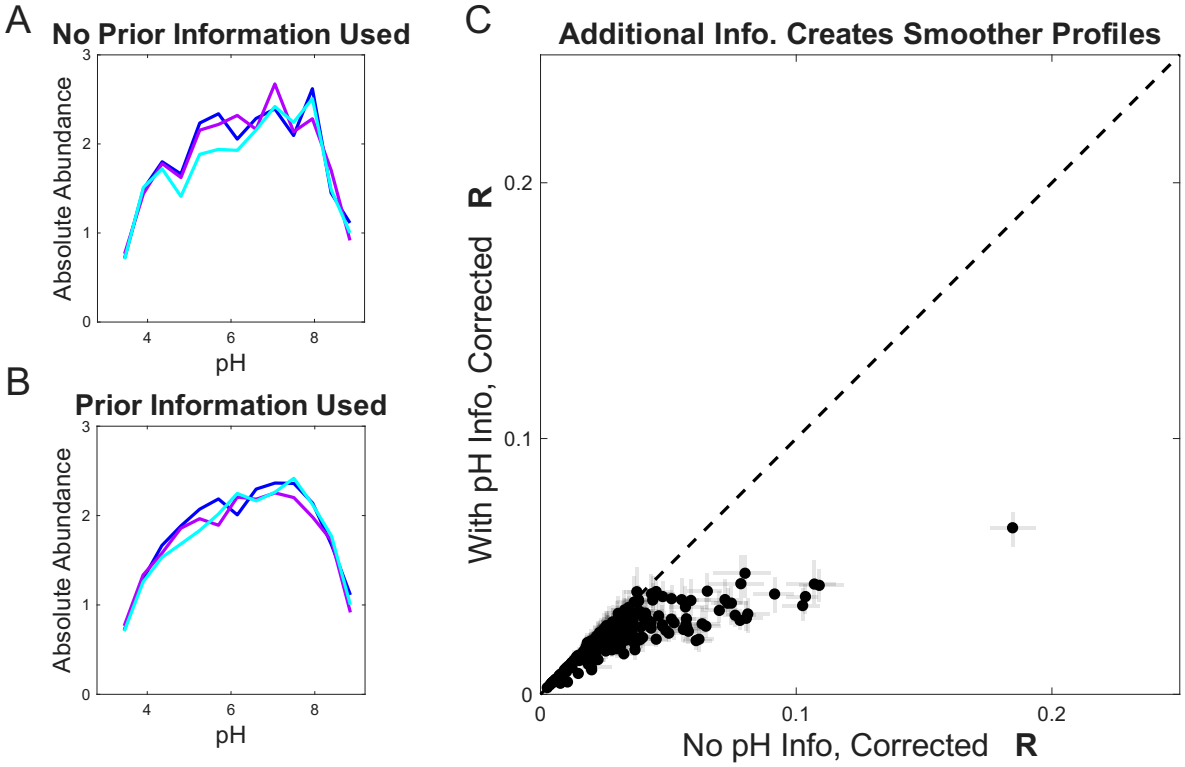
Prior expectations can be used to improve corrections. **A-B:** To compare the impact of using prior expectations, the profile of all 61 low-count sequences from Soil 17 in *Lee et al. (2025)* measured in all pH perturbations are shown with independent corrections applied with & without prior expectations. The prior expectation used was that the abundances should change smoothly as pH is varied (see **Methods**). When that prior expectation is used in the generation of corrections from training data, it also makes the example testing profiles smoother when compared to the profile generated with the standard corrections. **C:** To test the impact of including a prior expectation of smoothness with pH across all samples, corrections generated with & without the prior expectation were applied to an independent training set. **R** is the RMSE from a smoothened version of the data at each experimental condition (see **Methods**). Including a prior expectation of smoothness typically made testing data smoother. Error bars as in **Fig. 2A**.

## Discussion

It is uncontroversial to say that given a choice in the abstract between relative abundances or absolute abundances, absolute abundances would be preferred. All the information contained in relative abundances is also contained in absolute abundances. While the use of ISD’s allows for the indirect measurement of absolute abundances it also introduces a new source of systematic errors. This new source of systematic errors needs to be addressed for absolute abundances to deliver on their promise.

While existing approaches focus on modifying experimental design to reduce these errors (see [5], [6], [7] for examples), our work introduces a new method that can be generically applied to sequencing data to detect spike-in errors. The reliance of our method on pooling a large and diverse group of low-count taxa makes it best suited for high-diversity communities where such taxa are abundant [8, 9]. It is less suited for low-diversity settings like synthetic communities, where there may not be enough low-count taxa present in all technical replicates to generate accurate corrections. Our method also requires technical replicates—ideally three or more—since corrections are generated from deviations across shared replicates. Any sample with spike-in errors correlated to the sample conditions instead of individual replicates would be impossible for this method to correct (e.g. if a sample condition’s chemistry reduced the effectiveness of polymerase on the spike-in in all technical replicates of that sample). This same approach should also be applicable to sequencing data normalized by optical density, where errors would be caused by heterogeneous optical density between technical replicates.

All data analysis was performed using MATLAB (Mathworks, Inc.). The associated code and the subset of data from *Lee et al. (2025)* needed to reproduce all figures in this work are publicly available [10].

## Methods

### Noise model for sequencing with internal standards

In replicate *i*, the absolute abundance of a given sequence *µ* (*a*_*iµ*_) is determined by that sequence’s count (*c*_*iµ*_) and the replicate’s spike-in count (*s*_*i*_). We model the measurement of each sequence’s count as having a true value with stochastic noise of finite variance and average value zero (*ε*_*iµ*_ and *λ*_*i*_). Taxon *µ*’s measured absolute abundance in replicate *i* can then be written as

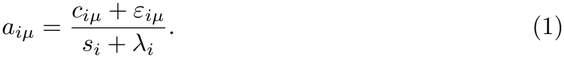

Assuming that the noise associated with the internal standard is small compared to the total spike-in count, we can approximate the measured abundance as

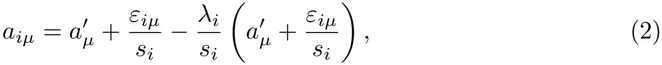

where *a*^*′*^ represents the true absolute abundance. We define *β*_*i*_ and *η*_*i*_ to be 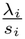and 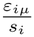, respectively. These are the noise terms measured in units of absolute abundance. While the magnitude of individual sequence counts and their noise could vary between replicates due to changes in sequencing depth, we assume their magnitude in absolute abundance will be similar between replicates. In practice this assumption is justified by using only low-count sequences, since if a sequence is low-count in all technical replicates any idiosyncratic variations must be small. Our method is based on combining many taxa into a larger group with abundance *A*:

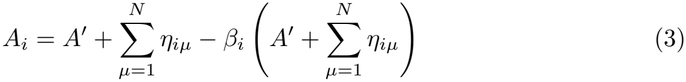

Keeping in mind that the true abundance *A*^*′*^ is constant, the variance of *A* across technical replicates is then

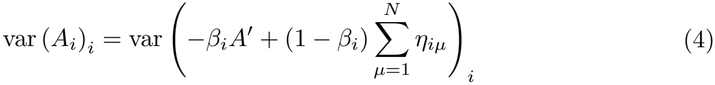

To simplify notation, future *i* indices will be dropped inside operators. This variance can be expanded to

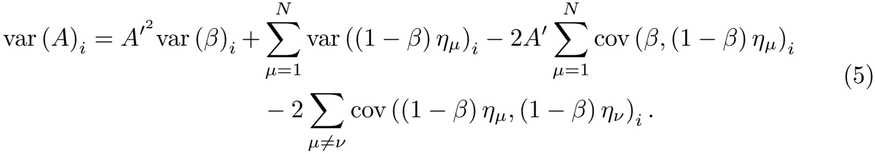

Assuming that noise sources for each sequence (including the spike-in) are independent and on average zero, the covariance terms are all zero. Note that for a given sequence *µ*,

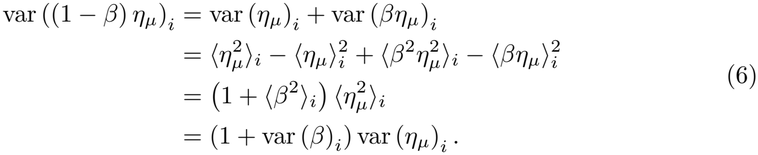

Using this identity to reduce var (*A*) further, we obtain:

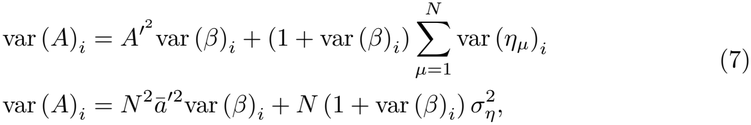

where 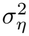 and *ā*^*′*^ are the average replicate-replicate variance of the combined sequences and the mean of the *N* absolute abundances being combined. As more sequences are added together, the variance of *A* becomes dominated by the internal standard error.

### Processing Data from *Lee et al*

The experiment in *Lee et al. (2025)* focused on measuring functional and compositional soil microbiome responses to pH perturbations. Experimental conditions where only one technical replicate existed were not used; this includes samples with the antifungal drug cycloheximide, as well as samples with both cycloheximide and the antibiotic chloramphenicol. Additionally, sample B44 in soil 15 was removed because only four sequences were identified in the sample. After filtering each sample to only include low-count sequences that were present in all technical replicates of a sample, the bottom 10% of samples with the fewest remaining sequences were discarded. This was done because the strength of a correction depends on the number of low-count sequences available to be combined, so samples where there are too few low-count sequences present in all technical replicates will be difficult to correct with this method. All other samples were used. In total, corrections were generated for 232 experimental conditions with 694 total samples.

### Internal Standard Correction Algorithm

Given a set of technical replicates, our algorithm aims to identify systematic errors attributable to errors in internal standard measurement. The algorithm can be summarized as:

1. Filter sequences
2. Bootstrap sequence groups
3. Calculate corrections
4. Generate consensus corrections

Specific details about each step and their implementation in this paper follow below.

#### Filtering Sequences

Any sequences not present in all technical replicates of a sample are removed. This maintains a consistent taxonomic composition for future steps that combine that sequence counts into larger groups. Next, sequences are filtered based on their abundance so that only low-count sequences are available for combining into groups. Under the assumption that *σ*_*η*_ is of similar scale to *a*, this constrains the size of *σ*_*η*_ to also be small. For this paper we used an average absolute abundance threshold of 0.01 for sequences to be considered low-count.

#### Bootstrap Sequence Groups

Given a set of *N* filtered sequences, the largest possible grouping is the combination of all *N* . Instead of relying on that single grouping, we generate an ensemble of groups of *N* sequences using bootstrap sampling to create multiple random combinations of sequences. Bootstrapping effectively tries a variety of weights for each sequence, which reduces the impact of any one sequence’s idiosyncratic variations on the final correction. Ideally, the combined group’s abundance is much larger than the abundances of any individual sequence and sequence noise. We generated 1000 coarsened groups via bootstrapping for the corrections studied in this paper. A correction is in error when it captures individual sequence variations rather than a systematic error. To estimate this uncertainty, the data a correction is trained on is repeatedly split in half, and two independent corrections are generated from each half. The average difference between those corrections is used as an estimate of the average impact of idiosyncratic variations on that correction. In this paper we generated error bars using 100 such splits of the training data.

### Calculating Corrections

Once we have an ensemble of random groupings, we generate corrections for each grouping based on its replicate-replicate variations. For a single grouping, corrections for each replicate are generated by calculating the factor that maps each replicate to an expected value for the true abundance of that group (*A*^*′*^ in **Eq. 3**). In this work we used the average value of the group across replicates for that expected value; however, better informed guesses could be used. For example, the pH informed corrections in **Fig. 3** assume that microbial abundances should vary smoothly with pH. This is incorporated by using, as the expected value, the average across the sample’s own replicates together with replicates from the neighboring pH conditions of the same soil. After generating corrections for each random grouping we have an ensemble of corrections for each replicate. We take the average across corrections to obtain a single consensus for each replicate.

## Acknowledgments

This work was supported in part by the Harvard Radcliffe Institute, by NSF grants PHY-2310746 and PHY-2340791, and by the National Institutes of Health under NIGMS Award Number R35GM160222. The content is solely the responsibility of the authors and does not necessarily represent the official views of the National Institutes of Health.

## Supporting Information

**Figure S1.**
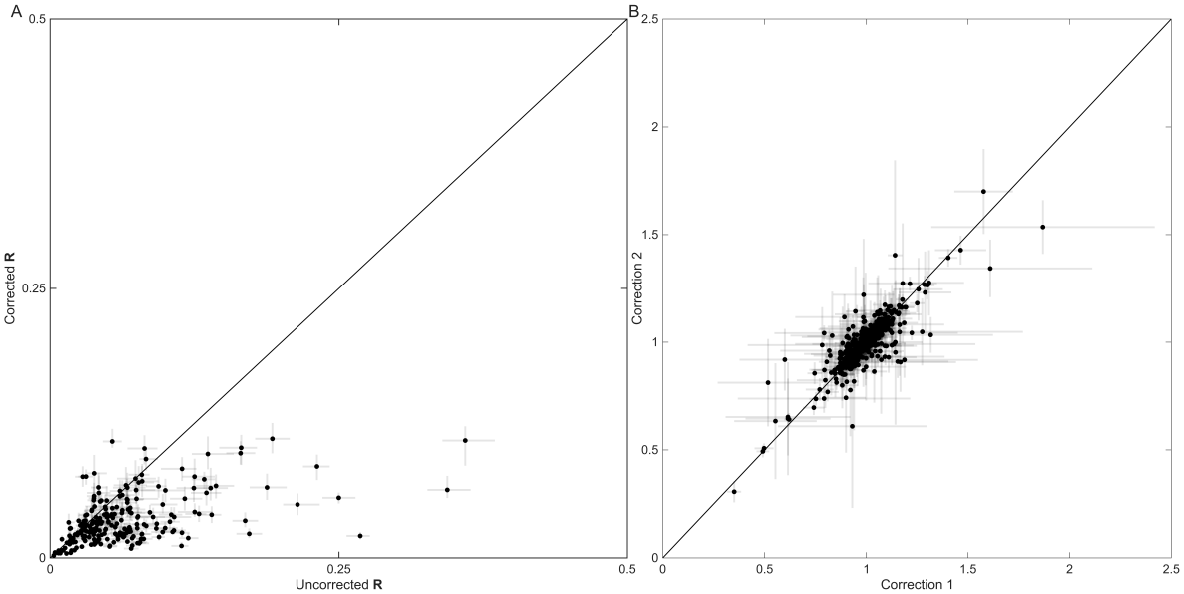
Generating corrections from large families. **A:** The same process as **Fig. 2** is applied to correct data, but instead of randomly grouping low-count sequences the 250 most abundant families are randomly grouped to generate corrections. Corrections are tested on random groupings of low-count sequences as in **Fig. 2**. In some cases the abundant families are able to be used to correct the spike-in error, but in some samples idiosyncratic noise or real biological variation is being corrected for instead. As a result, when corrections are applied to testing data they sometimes lead to increased variance. **B:** Corrections generated from two halves of the same training data in **A**. Idiosyncratic noise or actual biological variations make generating corrections dependent on the specific variations of each family. The increased sensitivity to the presence or absence of individual families in the bootstrapping portion of the correction pipeline leads to a high degree of uncertainty about the correction value, and often the value does not agree well with corrections generated from other subsets of the data.

**Figure S2.**
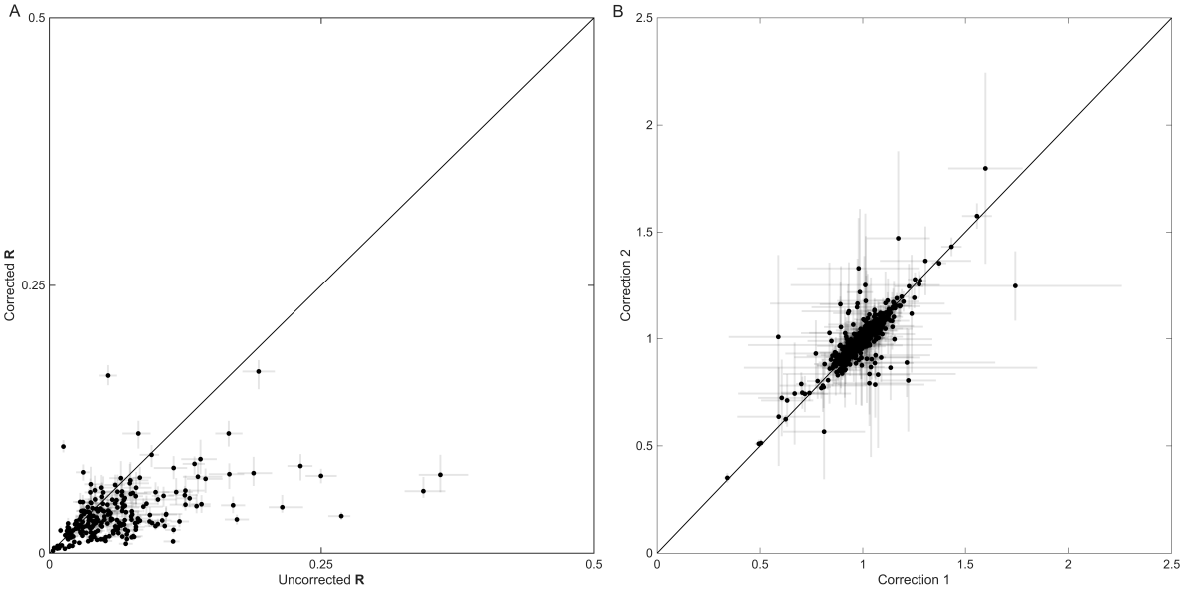
Generating corrections from high-count sequences. **A:** The same process as **Fig. 2** is applied to correct data, but instead of randomly grouping low-count sequences the 250 most abundant sequences are randomly grouped to generate corrections. Corrections are tested on random groupings of low-count sequences as in **Fig 2**. Again, highly abundant sequences are sometimes sensitive to spike-in errors, but sometimes they have their own variations that can be mistaken for spike-in errors. As a result, when corrections are applied to testing data they sometimes lead to increased variance. **B:** Corrections generated from different halves of the set of high abundance sequences. Idiosyncratic variations of individual sequences lead to large error bars for each correction and uncertain spike-in error detection.

**Figure S3.**
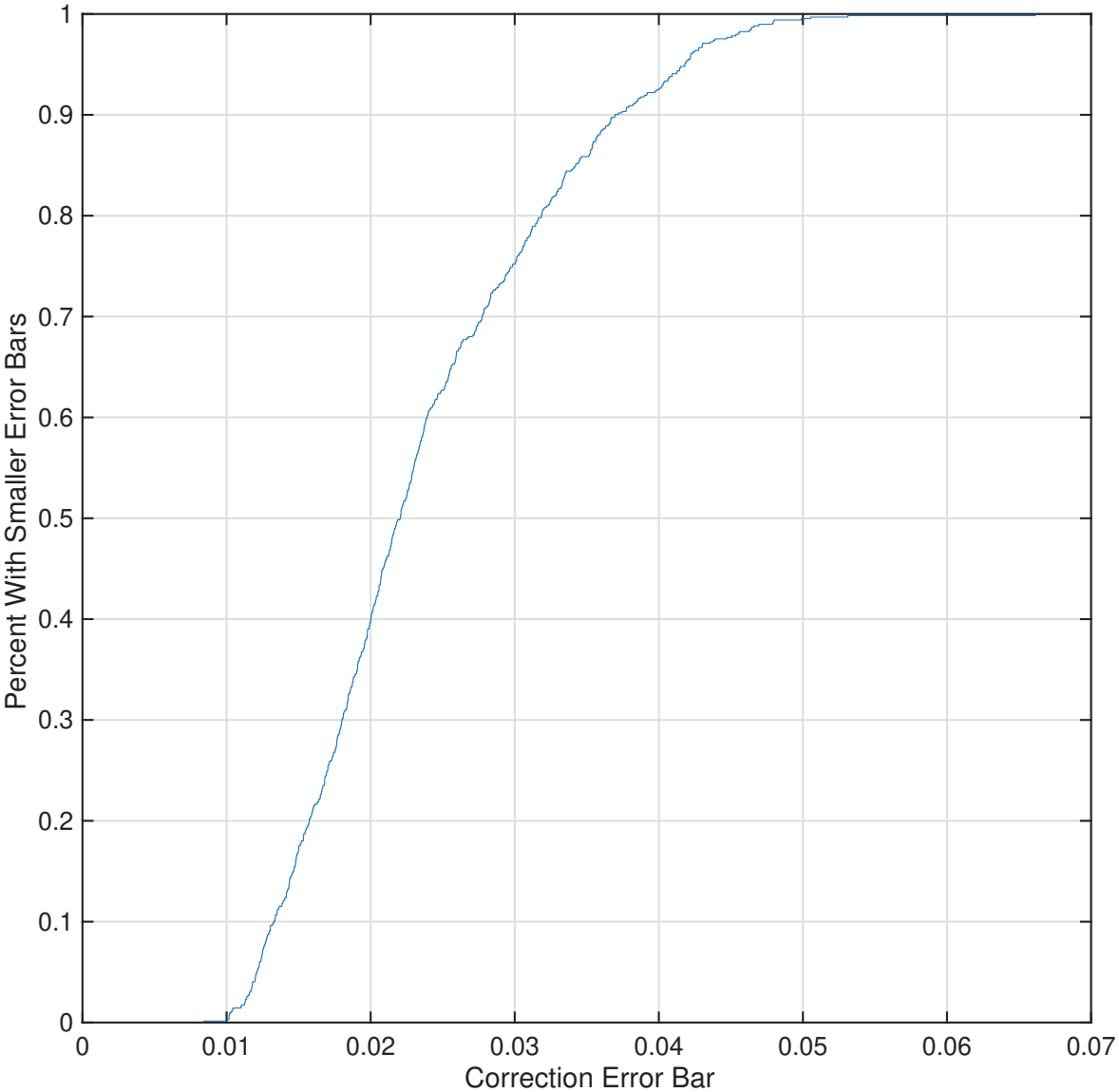
Cumulative distribution function of correction error bars. The corrections shown in **Fig. 2B** are generated from half of the training data; because they are generated from from smaller collections of low-count sequences they are more sensitive to individual sequences’ idiosyncratic variations and have larger error bars than corrections that would be generated with the entire available data. This cumulative distribution shows the percentiles of correction error-bars for corrections trained on the entirety of available data. 99% of corrections trained on the entire available data had error-bars smaller than 0.05.

